# A Spectrum of Time Horizons for Dopamine Signals

**DOI:** 10.1101/2021.10.31.466705

**Authors:** Wei Wei, Ali Mohebi, Joshua D. Berke

## Abstract

Dopamine input to striatum can encode reward prediction error, a critical signal for updating predictions of future rewards. However, it is unclear how this mechanism handles the need to make predictions, and provide feedback, over multiple time horizons: from seconds or less (if singing a song) to potentially hours or more (if hunting for food). Here we report that dopamine pulses in distinct striatal subregions convey reward prediction errors over distinct temporal scales. Dopamine dynamics systematically accelerated from ventral to dorsal-medial to dorsal-lateral striatum, in the tempo of their spontaneous fluctuations, their integration of prior rewards, and their discounting of future rewards. This spectrum of time scales for value computations can help achieve efficient learning and adaptive motivation for a wide range of behaviors.

## Introduction

How much should we care about the future? It makes sense to discount rewards that are far away in time - among other reasons, they are less certain to occur at all (1). Yet some worthwhile rewards take time and work to acquire. To maintain motivation and avoid choosing less favorable, but faster, gratification we must not discount delayed rewards too quickly. Excessive discounting - i.e., failure to maintain a sufficiently long time horizon - has been reported in a range of human psychiatric disorders (2), notably drug addiction (3).

The rate at which future rewards are discounted plays an important role in Reinforcement Learning (RL) theory, a widely-applied framework for understanding adaptive behavior in both animals and artificial agents (4). RL is built around discounted reward predictions (“values “). Values are updated using temporal-difference reward-prediction errors (RPEs) – mismatches between the value expected at each moment and ongoing experience. RPEs can be encoded by brief fluctuations in the firing of midbrain dopamine (DA) cells (5–9). DA cells project widely but especially to the striatum, a key brain node for value-guided decision-making (10, 11). RPE-scaled striatal DA release (12, 13) may engage synaptic plasticity (14, 15) to update values and subsequent behavior.

DA RPEs have been classically considered a widely-broadcast scalar signal (5). A single RPE signal implies a single underlying value, based on a single discount rate, and so defines a single time scale for learning and decisionmaking. However, animals typically need to make decisions, assess outcomes, and update their behavior accordingly over multiple time scales. During rapid production of motor sequences (e.g. birdsong) desirable (or not) results are produced by patterns of muscle activation a small fraction of a second before (16); it would be maladaptive to assign credit to actions performed much earlier. By contrast, other behaviors such as hunting for food can take orders of magnitude longer (1). Decisions to commit substantial time to an activity require a slow discount rate, and a correspondingly longer time window for updating values. Evaluation using multiple discount factors in parallel can better account for animal behavior (17, 18) and also improve performance of artificial learning systems (19, 20).

Furthermore, there is now substantial evidence for heterogeneity of DA cell firing (8, 21) and DA release across distinct striatal subregions (13, 22–28). These subregions are components of distinct large-scale loop circuits (29, 30), proposed to serve as distinct levels of a hierarchical RL architecture (31). Specifically, more dorsal/lateral striatal subregions are concerned with motoric details while more ventral/medial areas help to organize behavior over longer time scales (32). Theoretical studies have proposed a corresponding gradient of temporal discount factors across striatum (17) and there is some evidence for graded discounting from human fMRI (33). Yet how DA signals in distinct striatal subregions reflect distinct time scales for reward-related computations has not been examined, to our knowledge.

We report multiple lines of evidence for a gradient across the striatum of the time scales that determine dopamine dynamics. Most notably, we show that distinct subregions display very different patterns of dopamine release evoked by reward-predictive cues. We demonstrate that these patterns can be largely explained by distinct discount rates for underlying reward predictions, consistent with a portfolio of time horizons for decision-making.

## Results

### Dopamine tempo depends on striatal subregion

We used fiber photometry of the fluorescent DA sensor dLight1.3b (13, 35) to observe DA release fluctuations in the striatum of awake, unrestrained rats. We focused on three standard subregions (Fig. 1A; Extended Data Fig. 1): dorsolateral (DLS), dorsal-medial (DMS) and ventral (VS; targeting the Core of the nucleus accumbens). These receive distinct patterns of cortical input (36) and are often considered to have distinct “motor”, “cognitive” and “limbic” functions respectively (37, 38).

**Fig. 1.**
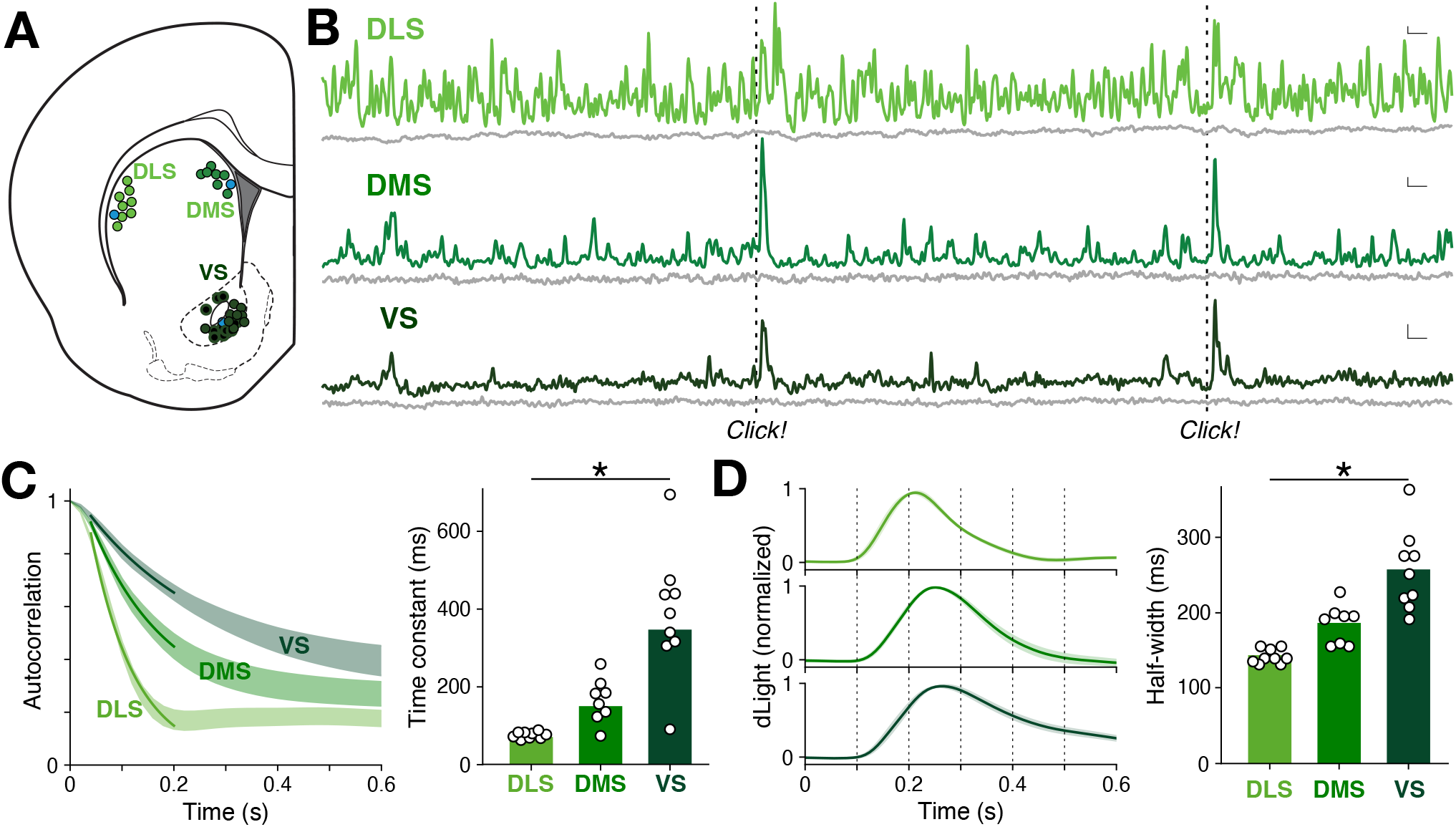
Dopamine tempo dependson striatal subregion. **A**,A rat brain atlas section (34), showing approximate locations of fiber optic tips (circles) within striatal subregions. Blue circles indicate the locations for the recordings in (B), and black-filled circles indicate locations for VS instrumental task recordings (13). For further details, see Extended Data Fig. 1. **B**, Example showing simultaneous, raw dLight photometry from each subregion in an awake unrestrained rat, outside of specific task performance. Green traces indicate DA signals (470 nm), grey traces indicate corresponding control signals (interleaved 415nm measurements). Occasional randomly-timed sugar pellet deliveries are marked as “Click!” (the familiar food hopper activation sound). Scale bars: 1s, 1% dF/F. **C**, Left, Average autocorrelogram functions for spontaneous dLight signals in each subregion. Bands show ± SEM, and darker lines indicate best-fit exponential decay for the range 40 ms to 200 ms. Data are from n = 13 rats over 15 recording sessions each; fiber placements n = 9 DLS, n = 8 DMS, n = 9 VS. Right, decay time constant depends on subregion (ANOVA: *F*(2,23) = 22.9, *p* = 3.4 × 10^-6^). **D**, Left, average dLight signal change after an unexpected reward click; right, duration (at half maximum) of signal increase depends on subregion (ANOVA: *F*(2,23) = 24.2, *p* = 2.2 × 10^-6^).

We first examined spontaneous DA fluctuations, unconstrained by task performance. DA dynamics were clearly different in each subregion (Fig. 1B; Supplementary Video). DLS signals showed near-constant, rapid change, while VS signals evolved more sporadically and slowly (Fig. 1C). When presented with a familiar, but unexpected, reward cue - the click of a hopper dispensing a sugar pellet - all three subregions showed a transient DA pulse (Fig. 1D). This pulse was briefest in DLS and lasted longest in VS (Fig. 1D). Prior voltammetry studies found that this same reward cue evoked DA selectively in VS (22), but our use of dLight may have revealed DLS/DMS responses that are too brief to readily detect with voltammetry. Briefer DA signals in more dorsal regions are consistent with studies showing faster rates of DA uptake, across species (39–41), although this alone appears insufficient to explain the highly distinct spontaneous DA events in simultaneous recordings (Fig. 1B).

### Distinct time scales for tracking reward history

As brief DA pulses can signal RPE, we next examined how the response to the reward click is affected by changing reward expectation, in each area. We took advantage of an instrumental task that we have extensively described before (13, 24). Well-trained rats made nose pokes, which sometimes produced the reward delivery click; reward probabilities shifted without warning between 10-90% (Extended Data Fig. 2). We previously reported that at reward delivery, both VTA DA cell firing and VS DA release scale with RPE - i.e. they are greater if fewer recent trials have been rewarded, reducing reward expectation. We now observed positive DA RPE coding also in DLS and DMS (Fig. 2A), although the DA pulse was briefer in DMS compared to VS, and again remarkably brief in DLS (Fig. 2B; half-width 121 ± 16 ms S.E.M.). On omission trials, DA dipped in all subregions, and the duration of this dip was also subregion-dependent (Fig. 2C).

**Fig. 2.**
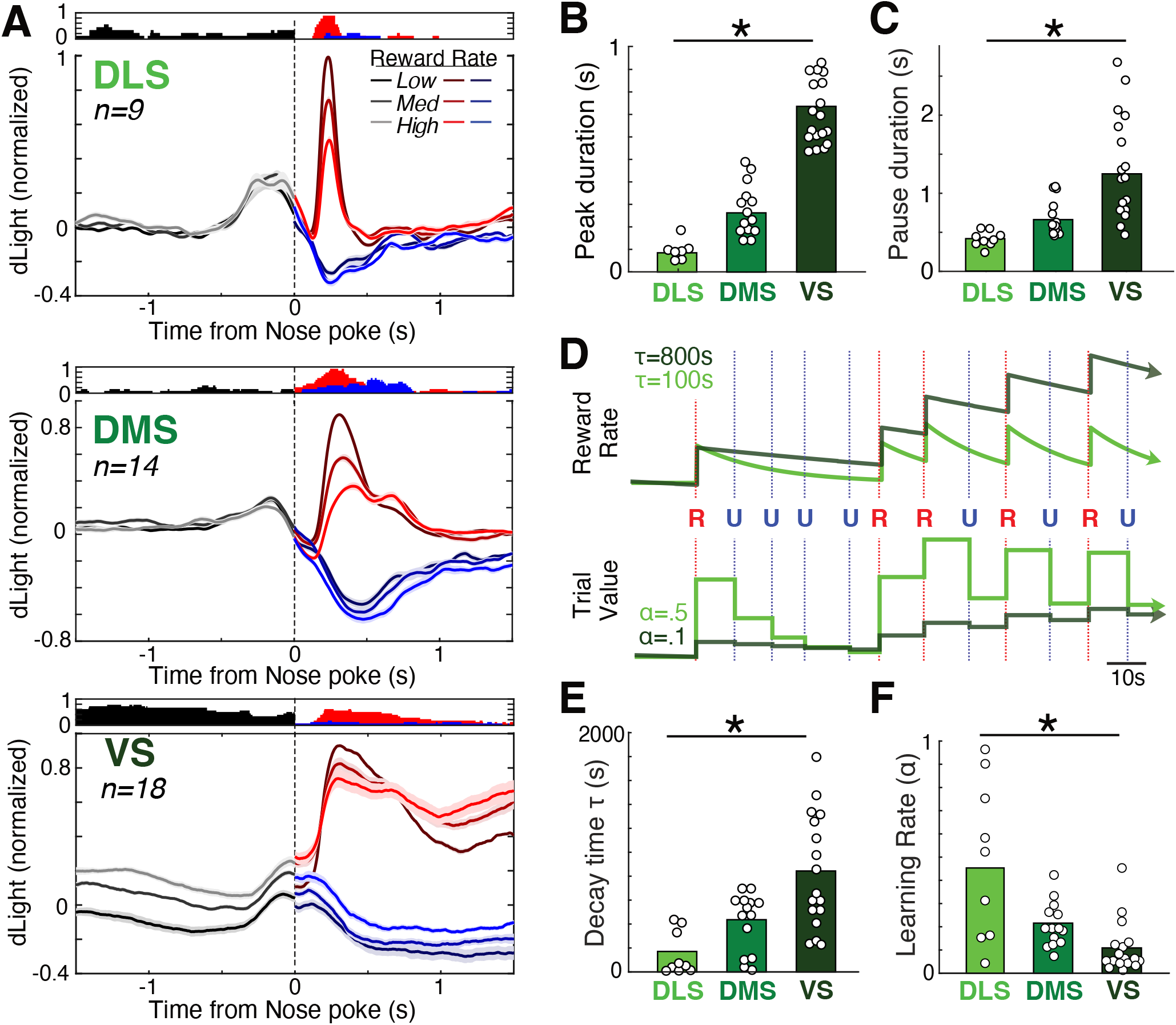
Dopamine prediction errors depend upon subregion-specific reward history timescales. **A**, Mean dLight DA signals aligned on reward delivery click, in the instrumental task. Data are from 12 rats, 1-3 sessions each; see Extended Data Fig. 1 for targets in each rat. Signals are broken down by recent reward rate (in terciles), with higher reward rate in brighter colors (error bars: SEM). After the time of nose poke, signals are further broken down by trials with (red) or without (blue) the reward click. Histogram above each plot shows the fraction of signals that significantly depended on reward rate (linear regression, *p* < 0.01), consistent with RPE coding after nose poke. Reward rates were calculated using a leaky integrator of reward receipts (see Methods and (D) below), choosing the *τ* parameter for each subregion separately to maximize RPE coding (alternative models of reward prediction or behavioral fits gave similar results, Extended Data Fig. 2). The bump before nose poke (most prominent for DLS) is the response to an earlier Go! cue, smeared by variability in reaction and movement times. **B**, The duration of the DA peak on rewarded trials significantly varies by subregion (repeated measures ANOVA, *F*(2,39) = 56.3, *p* = 4.4 × 10^-9^; measured at half-maximum in the 1s period after Side-In). **C**, same as (B) but for DA pause duration on unrewarded trials (repeated measures ANOVA, *F*(2,39) = 18.9, *p* = 9.4 × 10^-5^; half-minimum, 4s after Side-In). **D**, Top, illustration of leaky integrator estimation of reward rate, for an example sequence of trials (R = rewarded, U = unrewarded) and the *τ* decay parameter set to either 100 or 800s. Bottom, estimating reward expectation for the same example sequence using a simple delta-rule model, with one update per trial and learning rate parameter set to either 0.1 or 0.5. **E**, The leaky-integrator *τ* that maximizes correlation between RPE and DA after Side-In significantly varies by subregion (repeated measures ANOVA, *F*(2,39) = 23.6, *p* = 2.0 × 10^-5^). **F**, The deltarule learning rate *α* that maximizes correlation between RPE and DA after Side-In significantly varies by subregion (one-way ANOVA, *F*(2,39) = 23.2, *p* = 2.2 × 10^-5^). The strongest correlations are seen in DLS with a shorter time horizon (large *τ*, or small *α*) and in VS with a longer time horizon (small *τ*, or large *α*).

Despite being present in each subregion, the DA pulse was not a “global” RPE signal: it did not reflect the same underlying value in each subregion. Expectation of future reward can reflect past reward history over a range of possible (retrospective) time scales (42, 43). We estimated the time scale underlying each DA signal using a leaky integrator of rewards over time (44). This model has a single parameter *τ*: larger *τ* corresponds to a longer time scale, allowing rewards to better summate over multiple trials (Fig. 2D). For each DA signal, we determined the *τ* that produced the strongest correlation between DA pulses and RPE. We observed a systematic relationship to location: best-fit *τ* was shortest in DLS, intermediate in DMS, and longest in VS (Fig. 2E), consistent with a spectrum of time scales for reward history. This relationship to location was observed despite similar behavioral measures of reward expectation in the corresponding recording sessions (Extended Data Fig. 2).

As an alternative measure of the extent of recent history used to estimate rewards (7), we considered how quickly or slowly reward estimates are updated from trial-to-trial: smaller updates produce dependence on outcomes over a longer history of trials. Using a simple delta-rule model (45) we determined the learning rate *α* that maximized DA: RPE correlations at the reward click. Best-fit *α* was highest in DLS and lowest in VS (Fig. 2F), again indicating that VS is concerned with rewards integrated over more prolonged time scales.

### Region-specific responses to reward-predictive cues

We next turned from retrospective to prospective time scales for reward estimation. The RPE theory of DA function is based largely on DA cell responses to Pavlovian conditioned cues that predict future rewards (5, 9). Such responses are diminished when rewards are more distant, consistent with temporal discounting (46, 47). We examined DA cue responses in a Pavlovian approach task (Fig. 3A). Auditory cues (trains of 2, 5, or 9 kHz tone pips) predicted the reward delivery click a few seconds later, with distinct probabilities (75, 25, 0%; see Methods). Each trial presented one of the cues, or an uncued reward delivery, in random order, with a 15-30 s delay between trials. Rats were trained for 15 days, with 60 trials of each type per day. Early on, all cues increased the likelihood that rats would approach and enter the food hopper (Fig. 3B), consistent with generalization between cues (48). Over the course of training (3600 trials total), rats showed increasing discrimination, entering the food hopper in proportion to cued reward probability (Fig. 3B; Extended Data Fig. 3).

**Fig. 3.**
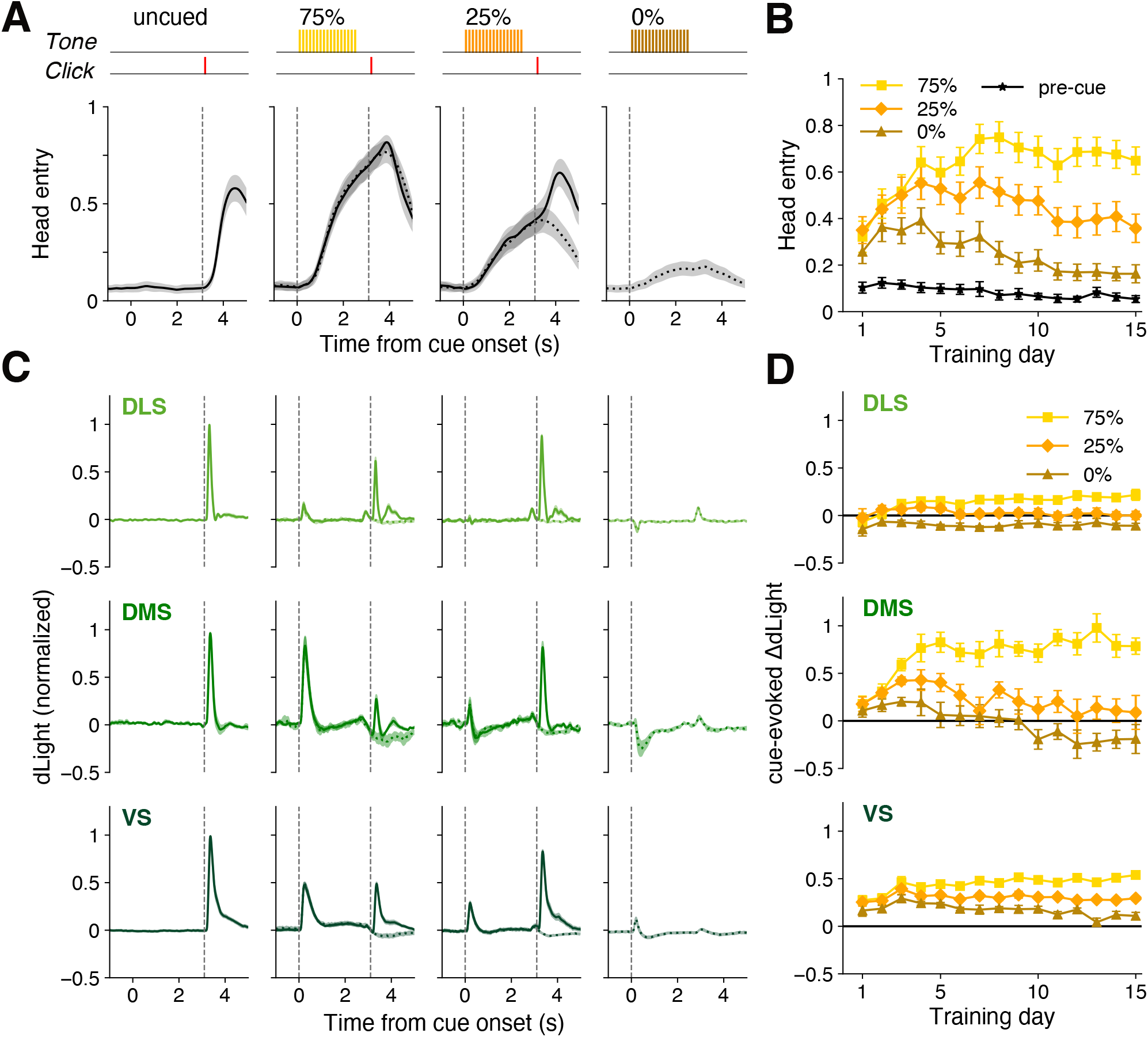
Subregion-specific dopamine responses to reward-predictive cues. **A**, Top, the Pavlovian task consists of four trial types, selected at random, with differing reward probabilities. Bottom, after training cues increase anticipatory head entries into the reward port (fraction of trials, mean ± SEM), and this scales with reward probability. Data shown are averages from training days 13-15, for n = 13 rats. **B**, During early training days rats increase their behavioral responses to all cues, before progressively learning to discriminate between cues (error bars: SEM; 2-way repeated measures ANOVA showed a significant CUE × DAY interaction (*F*(28,336) = 12.3, *p* < 10^-5^)). Points show average head entry in over a 0.5 s epoch just before cue onset (black) or just after cue offset (colors; i.e. immediately before the time that reward could be delivered). **C**, Average dLight signal change for each trial type after training (days 13-15, n = 13 rats with fibers in DLS (n = 9), DMS (n = 8) and VS (n = 9)). Solid lines = rewarded trials, dotted lines = unrewarded. **D**, Time course of DA increases to each cue in each subregion over training (mean ± SEM).By the late stage of training (days 13-15), the mean DA response depended on both cue identity and subregion (2-way ANOVA, significant CUE × AREA interaction, *F*(4,66) = 8.6, *p* < 0.0001). For more details on the development of behavior and DA responses, see Extended Data Figure 3.

These Pavlovian cues evoked strikingly-different DA responses in each subregion (Fig. 3C, D). By the end of training, DMS DA showed strong RPE coding: the 75% cue produced a strong DA pulse, the 25% cue a much smaller pulse, and the 0% cue a transient dip in DA (Fig. 3C). VS cue responses also scaled with RPE, but showed worse discrimination between cues, particularly on early training days, and remained positive for all cues throughout training (Fig 3D, Extended Data Fig. 3). Concordant results of VS DA increases to a learned 0% cue (CS-) have been previously observed and attributed to generalization between cues (49). Finally, in DLS all cues evoked much smaller DA responses (relative to unpredicted reward delivery). This did not simply reflect a failure of DLS-related circuits to learn: the DLS DA pulse at reward delivery was substantially diminished if preceded by the 75% cue (Fig. 3C), consistent with an acquired reward prediction.

### Weak DLS cue responses reflect very fast discounting

We reasoned that these subregional differences could reflect distinct time horizons for value computations. If future rewards are discounted especially fast in DLS-related circuits, even a brief delay would substantially diminish the value indicated by cues (Fig. 4A). To assess this, we turned to computational models that address the evolution of value within trials. We first applied a standard, very simple model in which the cue-reward interval is divided into a regular sequence of sub-states (the “complete serial compound”, CSC (50)). Over the course of learning, value propagates backwards along the sub-state chain (51). As expected, when we compared model versions with distinct discount rates (*γ*), rapid discounting reproduced the DLS pattern of smaller cue responses (Fig. 4B-D) despite a cue-dependent response to reward delivery (Fig. 4B). Including overlap between cue representations allowed the CSC to also reproduce generalization between cues early in training (Fig. 4D).

**Fig. 4.**
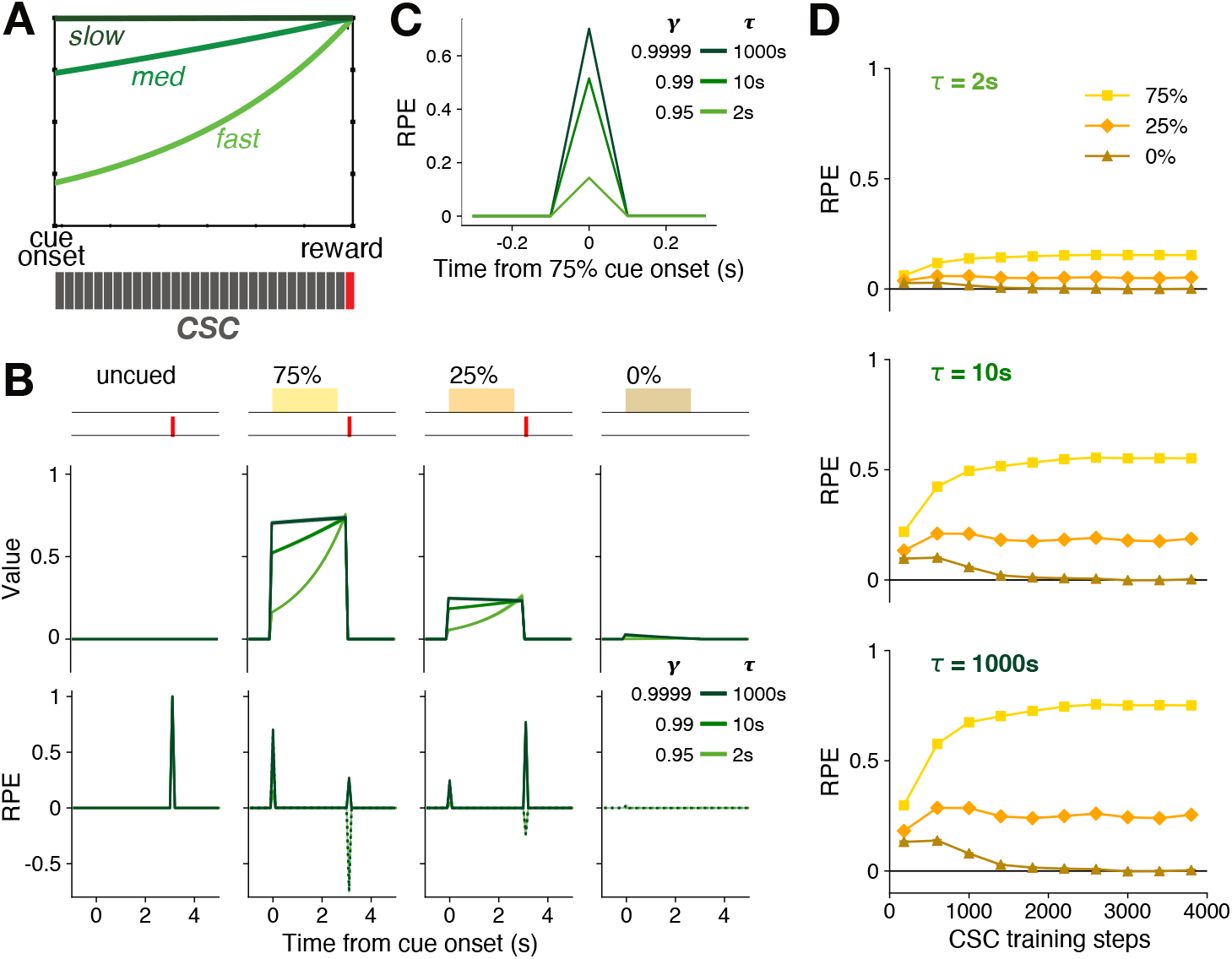
Faster temporal discounting can explain weaker DLS cue responses. **A**, Top, faster temporal discounting erodes the value indicated by the onset of a reward-predictive cue, even if the reward is certain to appear. Bottom, in the CSC model the cue-reward interval is divided into a fixed set of brief sub-states (we used 100ms duration). **B**, Values (top) and corresponding temporal-difference RPEs (bottom) for the CSC model after training in the Pavlovian task (step 3800). Discount factor *γ* was set to 0.95 (light green, “fast”), 0.99 (mid-green), or 0.9999 (dark green, “slow”). With a time step of 100 ms, these correspond to an exponential time constant (*τ*) of 2s, 10s, and 1000s respectively 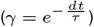. Even if the cued reward probability is high (75%), RPEs at cue onset are weaker when the discount factor is lower (RPEs at reward delivery are unchanged). **C**, Close-up of the CSC RPE response to the 75% cue. **D**, Development of RPEs at cue onsets with training. Note that cue discrimination is larger if *γ* is closer to 1 (plotted in more detail in Extended Data Fig. 4). Overlapping cue representations cause this CSC model to produce a positive RPE to the 0% cue early in training, but this fades to zero with extended training.

However, this CSC model of the cue-reward interval could not readily account for the slower, poorer cue discrimination in VS (Fig. 4C), and is incapable of reproducing the negative response to the 0% cue we saw in DMS. The model is not designed to handle prolonged time horizons that might span multiple trials (Fig. 5A; (52)). Furthermore, the splitting of experience into discrete, equally-fine sub-states becomes ever more artificial as inter-trial intervals get longer and more variable (53, 54).

**Fig. 5.**
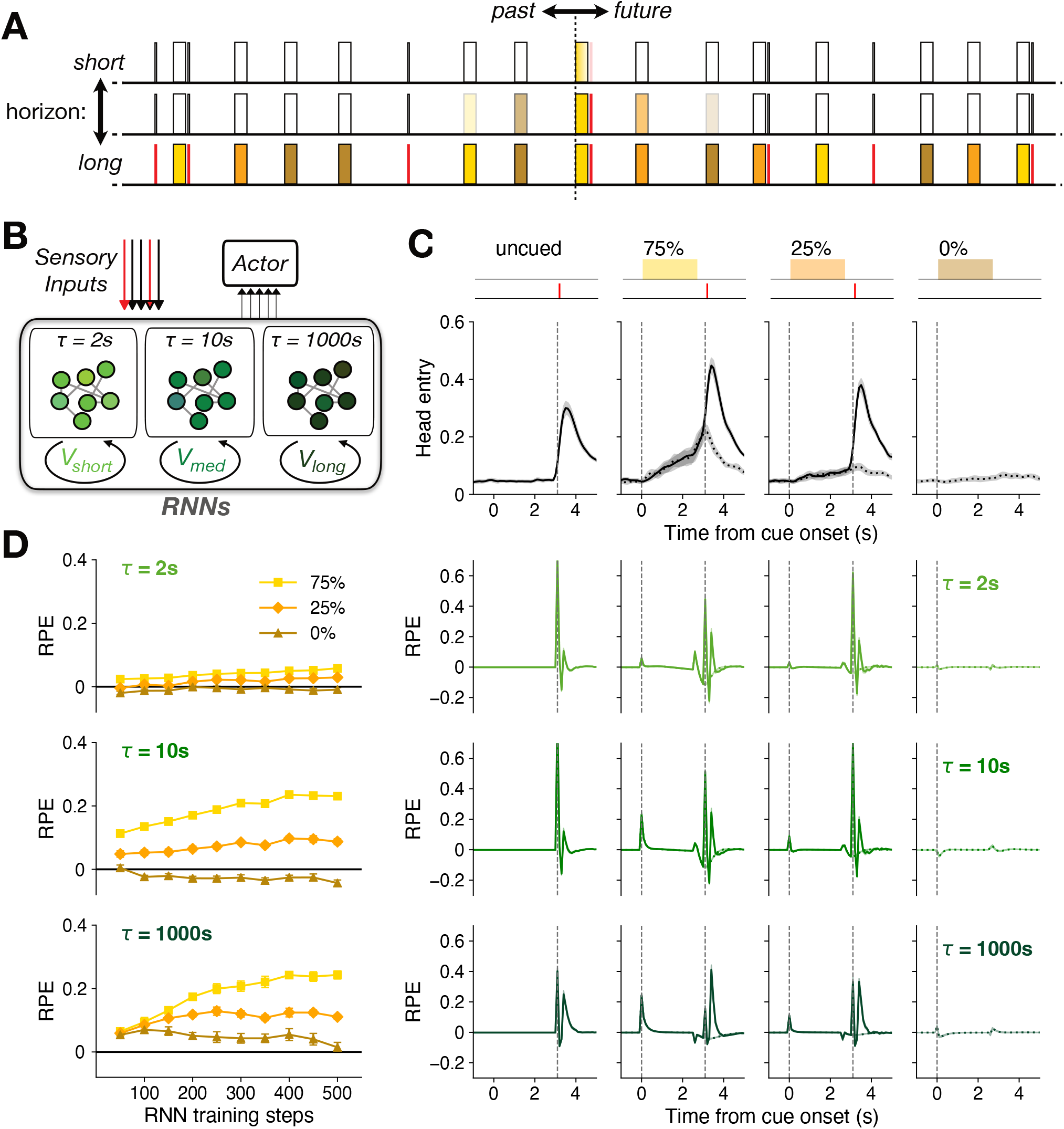
A longer time horizon accounts for slower VS cue discrimination. **A**, Schematic of part of a long random sequence of trials within a single training session, with colors indicating the cue in each trial. At any given moment, a reinforcement learning agent may be estimating the amount of reward that is coming “soon”, and updating such estimates based on what happened “recently”. If the time horizon is long, “soon” can encompass expected rewards across multiple trials, even if the current trial has 0% chance of reward. **B**, Schematic of recurrent neural network model, with three distinct pools of LSTM units. Each pool receives the same sensory inputs, but maintains its own value output based on a distinct discount factor (*γ* = 0.95, 0.99, or 0.9999, again corresponding to *τ* = 2s, 10s or 1000s). All three pools project to the Actor, which generates the probability of nose-poking. **C**, Model poke probability (top) and temporal-difference RPEs for each LSTM pool, after 500 training steps. **D**, Development of RPEs at cue onsets across training (see Extended Data Fig. 4 for extended training).

### Slow discounting impedes cue discrimination by VS DA

We therefore turned to an alternative approach for estimating the evolution of values, using recurrent neural networks (RNNs (55, 56)). In our composite RNN model (Fig. 5B; see Methods), each sub-network uses RL to generate distinct values in tandem (57), but with a distinct discount factor *γ* (58). The model has no discrete states and time is not explicitly represented, but rather is implicit within network population dynamics (59). With the sole assumption that *γ* increases from DLS to DMS to VS, the RPEs generated by the model recapitulated key distinct features of striatal DA pulses (Fig. 5C, D). These include the diminutive DLS responses as before, but also the negative DMS response to the 0% cue, and poor VS cue discrimination compared to DMS (especially earlier in training).

With extended RNN training, the “DLS” and “DMS” responses to cues remained relatively stable, but “VS” cue discrimination continued to improve, eventually also acquiring negative RPE responses to the 0% cue (Extended Data Fig. 4). In other words, a discount factor very close to 1 made learning slow, consistent with prior observations in RL models (60). With hindsight, this made intuitive sense. If the effective time horizon encompasses many trials, it will include multiple rewards regardless of which cue is presented on a given trial (Fig. 5A). Correctly assigning value to cues is therefore harder, and the discrimination is slower to learn. By contrast, if the time horizon for DMS is on the order of one trial, the average outcomes following distinct cues are very different (closer to the nominal 75, 25, 0%) and so learning the distinct associated values can be more quickly accom-plished.

The idea of distinct discount rates thus provides a concise explanation for the subregional differences in cue-evoked DA pulses. DLS responses are weaker because the cues indicate a reward that is too far away in time, given a short time horizon. VS responses are slower to discriminate, because the rewards that follow each cue are not very different, over a long time horizon. And DMS shows stronger, well-discriminating responses because its intermediate time horizon best matches the actual time scale of predictions provided by the Pavlovian cues.

## Discussion

A spectrum of time scales underlying DA RPEs is consistent with the hierarchical organization of behavior by cortical-basal ganglia circuits (31, 32, 61, 62). These time scales are also apparent in the representations and firing dynamics of individual neurons in these circuits (63, 64). DLS preferentially contributes to brief movements that can occur in rapid succession and require immediate feedback. This corresponds well to the rapidly-fluctuating DLS DA signal we observed in awake rats even outside of task performance (Fig. 1). DLS microcircuits have a range of features to support this faster tempo of information processing, including quicker DA reuptake and a higher proportion of fast-spiking interneurons to dictate fine timing (65). Operating on faster time scales with a more limited horizon complements the theory that DLS is involved in “habitual” stimulus-response (S-R) associations (38, 66). The key feature of S-R habits is that they do not take into consideration the future outcomes produced by actions – but in many behavioral situations, those outcomes may be simply too remote in time to alter DLS calculations.

Achieving adaptive behavior over longer time scales may require representations in VS circuits that are more prolonged and abstract (61, 67). Some imaging studies have suggested that VS circuits discount especially rapidly (33, 68) and may therefore promote maladaptive, impulsive behavior. By contrast, our results are consistent with an extensive literature demonstrating a critical role for VS in avoiding impulsive behavior (69, 70), by promoting work to obtain delayed rewards (71, 72). This slower discounting of future rewards is matched by a longer window for tracking past rewards (Fig. 2), as proposed by some theories of decision-making and time perception (73).

Our Pavlovian task used a standard systems neuroscience approach: cues that convey information about individual trials, with many trials in each session. But our results emphasize that animals, and their neural sub-circuits, do not necessarily process information in a corresponding trialbased manner (74). Slower discounting in VS may be important to motivate prolonged work, but can retard learning about cues that only provide information about the next few seconds. A VS time horizon that can span many trials may also explain puzzling observations of a large VS DA transient as each session begins (e.g., (75)). This makes sense if the onset of the first trial indicates that the animal is likely to receive multiple rewards “soon”, from the VS perspective.

Using multiple sub-agents with distinct discount factors may be a necessary strategy in a complex and changeable environment (43, 76). However, parallel cortical-basal ganglia circuits are not strictly segregated, but rather show convergence and connection (30, 77), consistent with overlapping information domains. This creates the challenge of how to appropriately integrate multiple, potentially-conflicting reward predictions (78). A multiplicity of discount rates has been previously proposed (17) to be responsible for choices that are inconsistent over time, a well-established feature of animal and human economic behavior (79, 80). An important question for future research is whether our increasing impatience as rewards draw near reflects the progressive engagement of more myopic dopamine-dependent valuation systems.

## ACKNOWLEDGEMENTS

We thank Nathaniel Daw, Vijay Namboodiri, Andrew Kayser, Robert Schmidt, and members of the Berke Laboratory for their comments on a prior version of the manuscript. Funding was provided by the National Institute on Drug Abuse (R01DA045783), the National Institute of Neurological Disorders and Stroke (R01NS123516, R01NS116626), the National Institute on Alcohol Abuse and Alcoholism (R21AA027157), the National Institute on Mental Health (K01MH126223), and the UCSF Wheeler Center for the Neurobiology of Addiction.

## AUTHOR CONTRIBUTIONS

A.M. performed the behavioral photometry experiments and instrumental task analyses. W.W. performed the computational modeling and Pavlovian task analyses. J.B. developed the conceptual framework, oversaw the study, and wrote the manuscript.

## Methods

### Animals and Behavior

All animal procedures were approved by the University of California, San Francisco Animal Care Committee. N=20 adult wild-type Long-Evans rats (15 males) were bred in-house, maintained on a reverse 12:12 light: dark cycle and tested during the dark phase. All recordings were performed in an operant chamber (Med Associates), and details on both behavioral tasks have been published previously (13, 24). For the Pavlovian task each cue tone (2, 5 or 9 kHz) was presented as a train of pips (100 ms on, 50 ms off) for a total duration of 2.6 s followed by a delay period of 500 ms. Trials with one of the three cues, or an unpredicted reward delivery, were delivered in pseudorandom order with a variable inter-trial interval (15–30 s, uniform distribution). Instrumental task sessions used the following parameters: left–right reward probabilities were (independently-varying, randomly-selected) 10, 50 or 90% for blocks of 35-45 trials; hold period before the Go cue was 500–1,500 ms (uniform distribution). The mean number of trials for included recording sessions was 300 (range: 164-407).

### Virus and Photometry

We used a viral approach to express the genetically-encoded optical DA sensor dLight1.3b (35). Under isoflurane anesthesia, 1 μl of AAV-DJ-CAG-dLight1.3b (2 × 10^12^ viral genomes per ml; Vigene) was slowly (100 nl/min) injected (Nanoject III, Drummond) through a glass micropipette targeting multiple striatal subregions: ventral (AP: 1.7, ML: 1.7, DV: 7.0 mm relative to bregma), dorsomedial (AP: 1.5, ML: 1.8, DV: −4.3) and dorsolateral (AP: 0.84, ML: 3.8, DV: −4.0). During the same surgery optical fibers (400 μm core, 430 μm total diameter) attached to a metal ferrule (Doric) were inserted (target depth 200 μm higher than AAV) and cemented in place. Data were collected >3 weeks later, to allow for dLight expression. For dLight excitation blue (470 nm) and violet (405 nm; isosbestic control) LEDs were alternately switched on and off in 10ms frames: 4ms on and 6ms off (81). Excitation power at the fiber tip was set to 30 μW for each wavelength. Both excitation and emission signals passed through minicube filters (Doric) and bulk fluorescence was measured with a femtowatt detector (Newport, Model 2151) sampling at 10 kHz. Time-division multiplexing produced separate 470 nm (DA) and 405 nm (control) signals, which were then rescaled to each other via a least-square fit (82). For the simultaneous recording of three areas, we used a Neurophotometrics system (83); technical details were very similar except that the control wavelength was 415nm and detection was camera-based, sampling at 100 Hz. Fractional fluorescence signal (dF/F) was then defined as 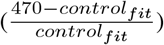. For each Pavlovian recording session DA activity was normalized to the mean peak uncued click response in that session. We removed from analyses 3 fiber placements that produced consistently weak signals (1 DMS, 2 VS), and we also excluded individual sessions for which the peak response was less than one standard deviation (Z < 1; 19 of 390 sessions excluded, 2 DLS, 15 DMS, 2 VS). DA activity at cue time was estimated as the maximum or the minimum within a half second window after cue onset, whichever had the larger absolute value; results were not substantially different if we instead used average DA in this window (data not shown).

### Histological confirmation

To verify probe placement post-mortem, animals were perfused transcardially with PBS and then 4% PFA. Implants were taken out and brains were extracted and postfixed in 4% PFA for 24 h, then placed in 30% sucrose in PBS for >48 h and sectioned at a 100μm thickness with a microtome. We used immunofluorescence staining to visualize dLight expression. Brain sections with probe placement were identified, blocked in a 0.4% Triton X-100 solution with 5% normal goat serum (NGS) for 1 h at room temperature, followed by an overnight incubation in a rabbit anti-GFP primary antibody solution (1:1000; abcam, ab290) in PBS in a cold room. Sections were washed three times in PBS for 10 min at room temperature and incubated in an Alexa 488-conjugated goat anti-rabbit secondary antibody solution (1:250) in PBS for 1 h at room temperature. Finally, sections were washed six times in PBS for 5 minutes at room temperature and then mounted onto glass slides and coverslipped using Fluoromount-G™ Mounting Medium, with DAPI. Fluorescent images were taken using a fluorescence microscope (Keyence BZ-X810) with a 2x objective lens. Fiber tip locations from both hemispheres were projected onto the same side in atlas space.

### Computational Models

#### Trial-level models

For the instrumental task we estimated reward rate using a time-based leaky-integrator. Reward rate was incremented by 1 at each time the rat received a reward, and exponentially decayed with time constant *τ*. *τ* was varied between 1-2500s, to find the strongest negative correlation between reward rate and the DA peak after Side-In (within 0-1s, on rewarded trials; i.e. positive RPE coding). To estimate learning rate, we used a trial-based delta-rule. This model tracks a state value that is updated once per trial by *V*(*t*) = *V*(*t* – 1) + *α* * (*r* – *V*(*t* – 1)); *V*(*t*) is the trial-based state value at trial t, *α* is the learning rate and r is the outcome of each trial (0 or 1). By varying the value of *α* between 0 and 1 (in 0.01 steps) we found an optimal value for each DA signal that would minimize the correlation between state value and peak DA signal in a 1s window after Side-in.

#### Real-time models

The CSC model is a standard temporal-difference model of conditioning (50). Values are defined as a linear function of features *x* and weights *w*, 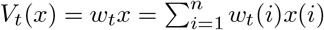, where n is the time steps in a trial. The vector *x* is non-zero only at the *t_th_* element at time step t after cue onset, i.e., *x*(*i*) = *δ_it_*, where *δ_it_* is the Kronecker delta function. In addition to activating a single distinct feature for each cue, we also included shared features activated by any of the three cues, to allow for generalization. In the results presented we used a single shared feature, but increasing the number of shared features did not qualitatively affect results (not shown). The weights *w* update according to *w*_*t*+1_ = *w_t_* + *αδ_t_e_t_*, where *α* is the learning rate (we used *α* = 0.01), *δ_t_* is the RPE and *e_t_* is an eligibility trace. The RPE is defined as *δ_t_* = *r_t_* + *γV_t_*(*x_t_*) – *V_t_*(*x*_*t*–1_), where *γ* is the discounting factor. The eligibility trace *e_t_* is included to accelerate learning and updated by *e*_*t*+1_ = *γλe_t_* + *x_t_*, where *λ* is a decay factor (we used *λ* = 0.98). The CSC model was run separately for each discount factor.

The RNN model, based on an advantage actor-critic architecture (84), is composed of LSTM units (85). These are organized as three sub-networks (“DLS”, “DMS”, “VS”) of 32 nodes each, with internal recurrent connections but without direct connections between sub-networks. Each sub-network receives the same copy of the sensory inputs at each time point, and generates its own value estimate using a distinct discount factor. All three sub-networks project to the same policy component, together generating the probability for taking an action (either “poke” or “no-poke”). These probabilities are sampled to determine the action at each time step. We used a time step of 100 ms.

The vector of sensory inputs to the RNN, include the food delivery click (0 for no-click or 5 for click), auditory cues, and background dimensions. Background dimensions (3d, all set constantly to 1) are included to mimic the background or contextual inputs to the network. The auditory cues consist of 20 dimensions, of which 3d are the distinctive one-hot features of the three cues and the remainder are set to 1 during all cue presentations to produce similarity between cues.

At each time step the RNN model receives reward feedback. Before reward delivery, the reward is 0 for taking the action “no-poke”, and −0.006 for taking the action “poke”, i.e., there is a small poking cost to discourage constant poking. If the poke output is maintained on consecutive time steps, the cost is reduced to 10% of that for first poke. In a rewarded trial, the reward (with value 1) is presented at a delay of 3 time steps after the reward delivery click.

The network was trained to perform the conditioning task by minimizing a loss function with three terms,

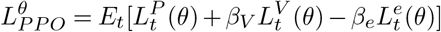

where the expectation was over a sequence of time steps with length T. We used T = 5000 steps, which encompasses multiple (~ 20) trials. We took the proximal policy optimization (PPO) for estimating the policy loss, which has the following form (86)

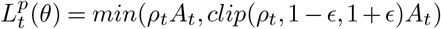

where 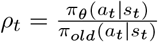 is the probability ratio, whose value is clipped with a parameter *ϵ*. The advantage *A_t_* includes three components,

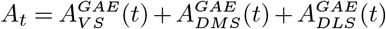

where each term is the generalized advantage estimator (GAE) (87) from one of the three sub-networks. Take the VS term as an example and define 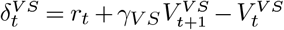 as the RPE at time t, then

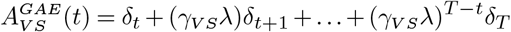

where T is the sequence length and *λ* is a parameter for GAE.

The value loss was given by

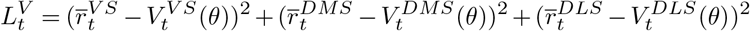

where, 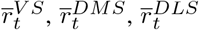 are the expected discounted rewards within the sequence, given the corresponding discount factor for each subnetwork. We used the value right after T to bootstrap the contribution from rewards beyond this sequence. For instance, the expected reward for VS has the following expression

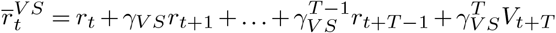

Since *γ_VS_* is very close to 1, the expected reward for “VS” sub-network reflects contributions from multiple trials. Faster discounting for “DMS” and (especially) “DLS” sub-networks results in minimal contributions from subsequent trials. The entropy term *L^e^* represents the entropy of the probability distribution of taking the two actions and was added to encourage the exploration. The parameters used were: *β_V_* = 0.8, *β_e_* = 0.001, *γ_VS_* = 0.9999, *γ_DMS_* = 0.99, *γ_DLS_* = 0.95, *λ* = 0.98. The weights of the network were updated using the Adam method (88), with learning rate 0.0005.

---

**Extended Data Figure 1.**
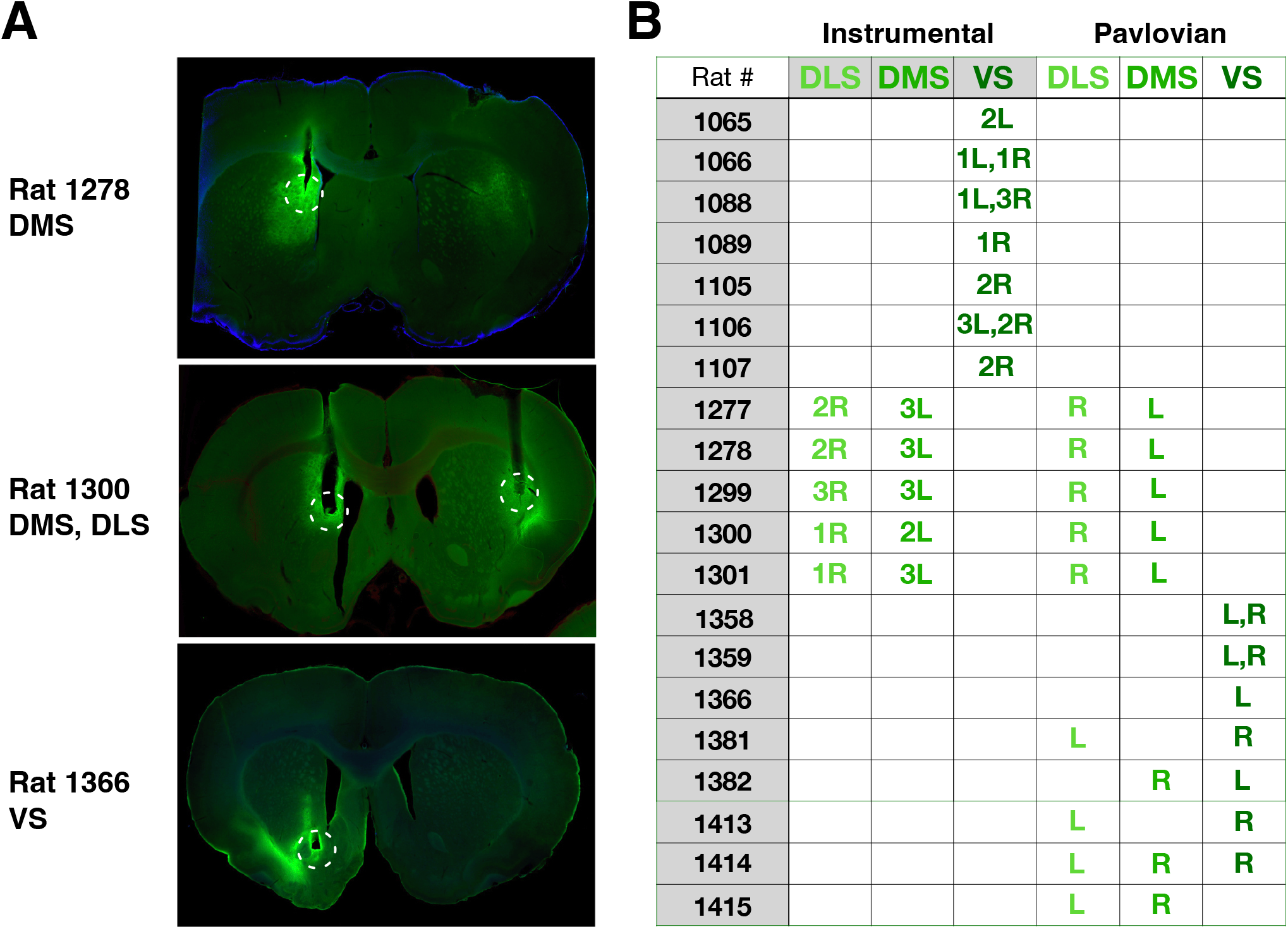
Photometry recording locations. A, Histology examples showing optic fiber tip locations (circled) and dLight1.3b expression (green), in DLS, DMS, VS. B, Table showing included fiber subregions for each rat and task. “L” indicates left hemisphere, “R” indicates right. For the instrumental task, numbers (1–3) indicate that multiple sessions were included for that fiber placement. A subset of data from rats 1065-1107 were previously reported (13).

**Extended Data Figure 2.**
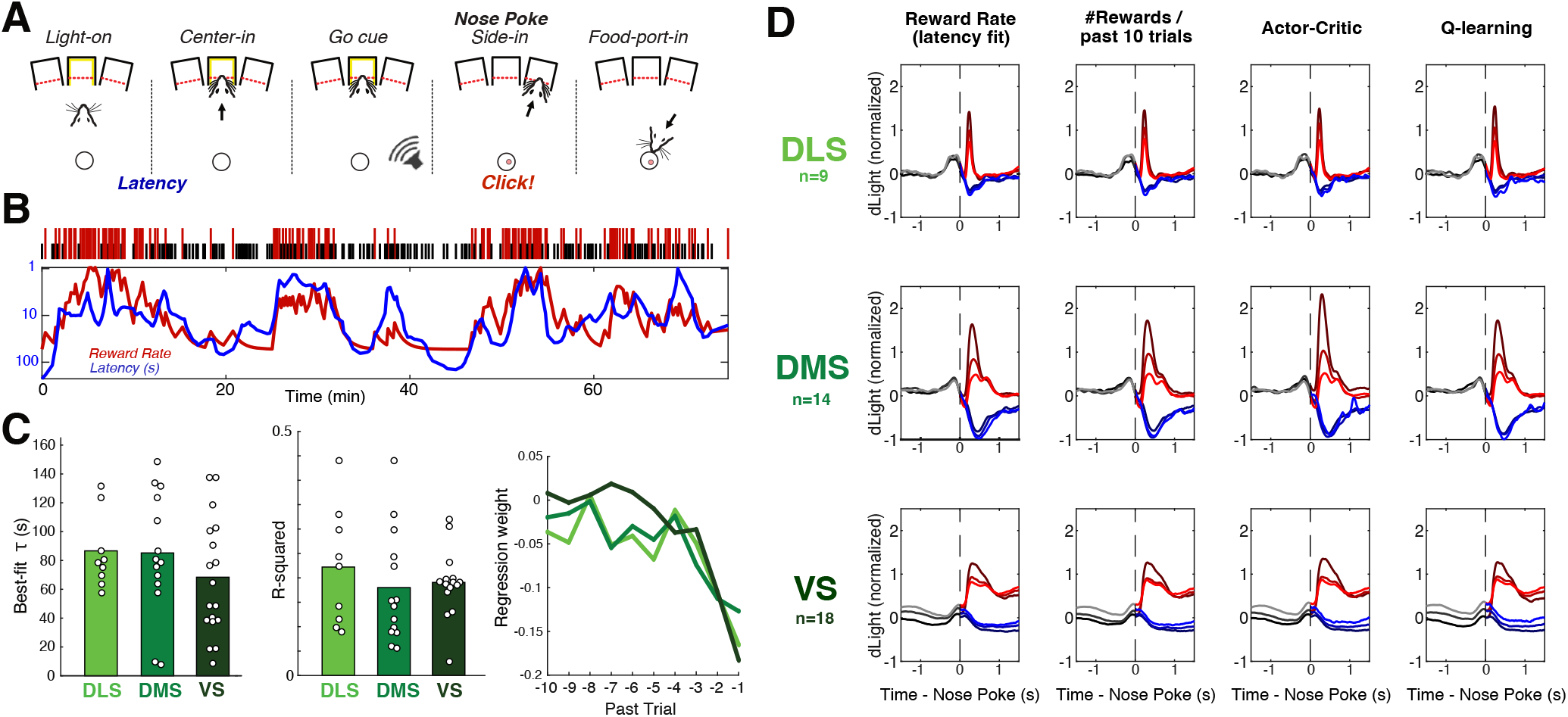
Instrumental behavior and alternative RPE fits. **A**, Schematic of instrumental task events. Here we focus on DA signals following the nose poke at Side-in, when the rat discovers if the current trial will be rewarded (food hopper click) or not (for information about other events see (13, 24)). As a measure of reward expectation we use “latency” (the time between initial Light On and the rat’s Center-in nose poke. **B**, Example behavioral session showing fit between latency (log scale, inverted) and recent reward rate. Tick marks at top show the timing and outcome of each trial (taller red ticks indicate rewarded trials, shorter black ticks unrewarded). Graphs show latency (5-trial running average) and reward rate, calculated with a leaky integrator using the *τ* parameter that produced the strongest (negative) correlation between latency and reward rate. **C**, Left, best-fit *τ* for each session in which DLS, DMS, and/or VS signals were recorded. There was no significant behavioral difference between recording locations (repeated measures ANOVA, *F*(2,39) = 1.72, *p* = 0.197). Middle, the amount of variance in latency that was explained by best-fit reward rate did not differ by recording location (repeated measures ANOVA, *F*(2,39) = 0.180, *p* = 0.673). Right, Coefficients of multiple regression examining effects of the outcome of the preceding 10 trials on (log) latency, separately for each subregion (same colors as bar charts). **D**, Alternative estimates of reward expectation produce similar RPE results. Each column uses the same data and format as Fig. 2A. From left, “Reward Rate” is also based on a leaky integrator, but using the *τ* best-fit to latency (as in B/C). “Rewards in the past 10 trials” is a simple count. “Actor-Critic” uses the Critic value from a trial-based Actor-Critic model, fitting the Critic learning rate to behavioral latency and the Actor *α*, *β* parameters to left and right choices. Q-learning uses a trial-based Q-value model, fitting the *α* and *β* parameters to choices and using Q (chosen action) as reward expectation.

**Extended Data Figure 3.**
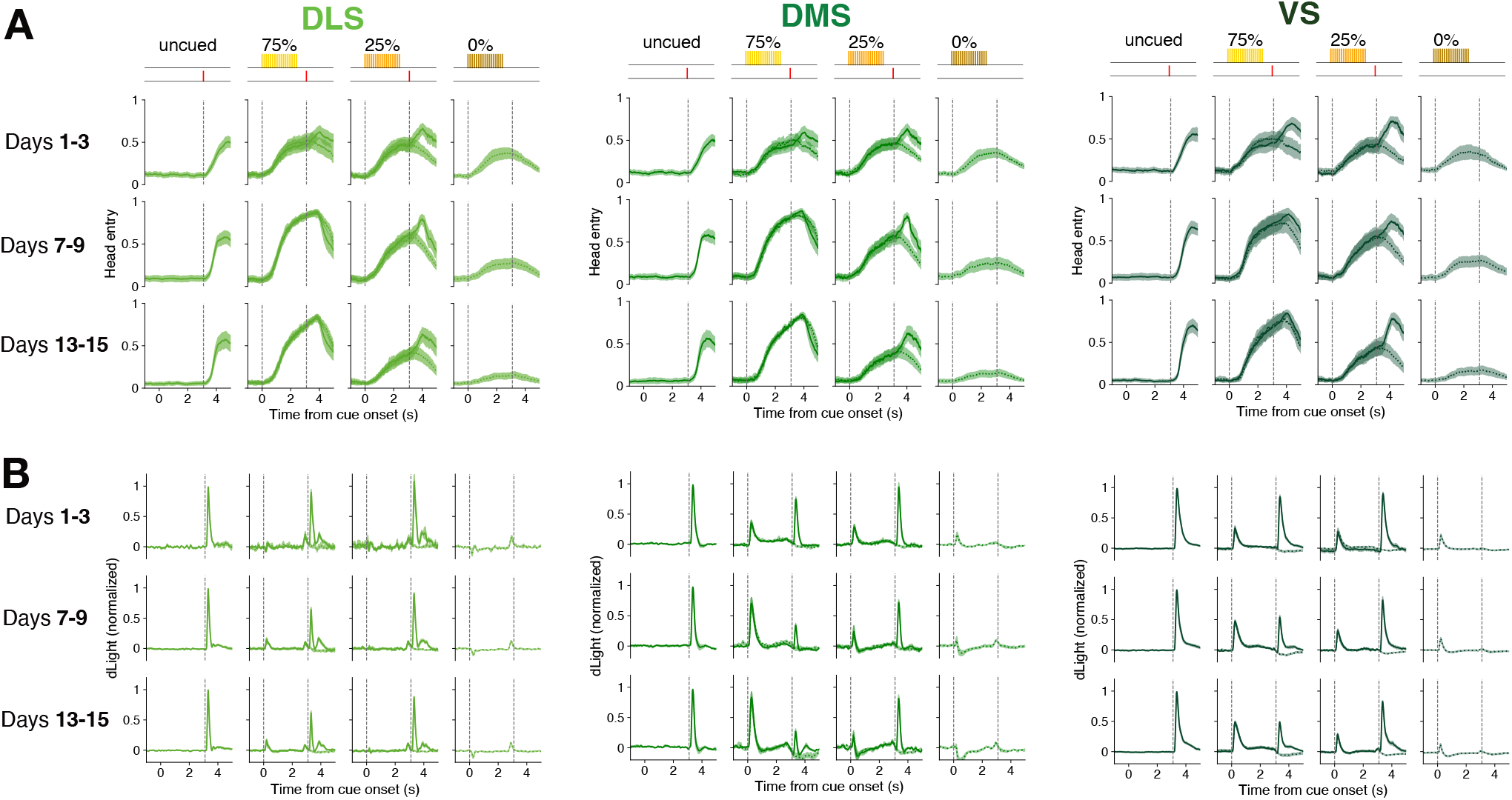
Development of approach behavior and DA cue responses in each subregion. **A**, Head-entry behavior develops in a very similar way regardless of recording site. Data shown is averaged across days 1-3, 7-9 or 13-15 respectively. **B**, Same sessions as A, but showing mean DA responses during each trial type. In all subregions discrimination between cues increases with time, but this is slow in VS.

**Extended Data Figure 4.**
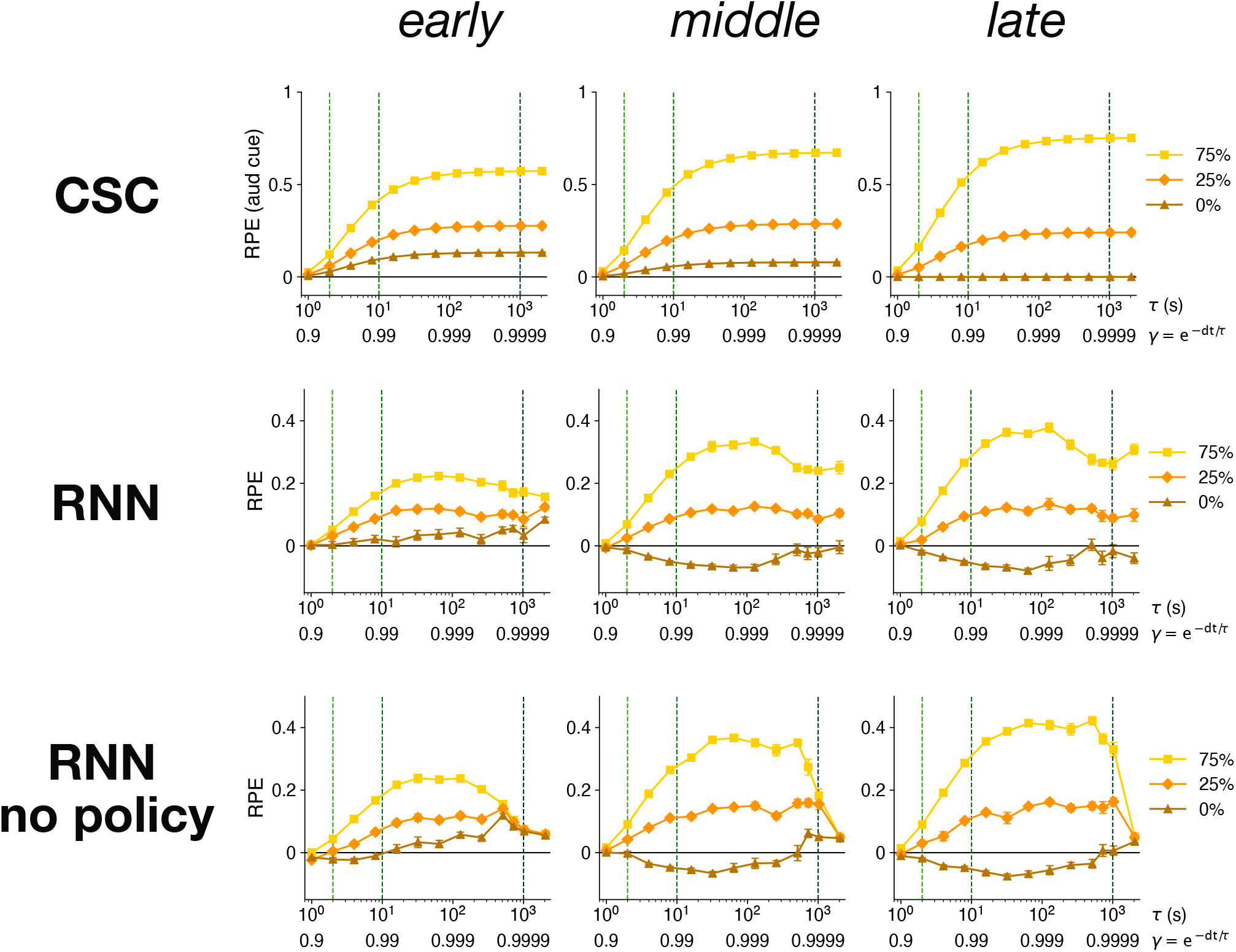
Effects of extended model training on cue discrimination with different discount factors. Top row, cue-evoked RPEs in the CSC model at “early” (600 training steps), “middle” (1000) and “late” (3800) stages of learning, as a function of *γ*, or equivalently the time parameter *τ*. (*γ* = *e^-dt/τ^*, where dt is the time step size, here 100ms). Green dashed lines mark *γ* = 0.95, 0.99, and 0.9999. Note that for low *γ* all cue responses are small even after learning, since any potential reward is heavily discounted. This CSC model initially shows a positive response to the 0% cue due to overlapping cue representations; over training this response fades to zero (but cannot become negative). Middle row, same for an RNN model (early = 100, middle = 500, late = 900 training steps). To isolate the effect of varying *γ*, this model variant used just a single network (a single *γ*) rather than three. Note that at early and middle stages of learning, if *γ* is close to 1 the RNN model shows less discrimination between cues compared to intermediate *γ*, consistent with the observed difference between VS and DMS. Bottom row, same as middle row, but also removing the Actor (poking) component.

